# Orthogonal genome-wide screenings in bat cells identify MTHFD1 as a target of broad antiviral therapy

**DOI:** 10.1101/2020.03.29.014209

**Authors:** Danielle E Anderson, Jin Cui, Qian Ye, Baoying Huang, Wenhong Zu, Jing Gong, Weiqiang Liu, So Young Kim, Biao Guo Yan, Kristmundur Sigmundsson, Xiao Fang Lim, Fei Ye, Peihua Niu, Xuming Zhou, Wenjie Tan, Lin-Fa Wang, Xu Tan

**Author notes:** These authors contributed equally to the work (Danielle E Anderson, Jin Cui, Qian Ye, Baoying Huang). To whom correspondence should be addressed: Xu Tan or Lin-fa Wang.

## Abstract

Bats are responsible for the zoonotic transmission of several major viral diseases including the 2003 SARS outbreak and the ongoing COVID-19 pandemic. While bat genomic sequencing studies have revealed characteristic adaptations of the innate immune system, functional genomic studies are urgently needed to provide a foundation for the molecular dissection of the tolerance of viral infections in bats. Here we report the establishment and screening of genome-wide RNAi library and CRISPR library for the model megabat, *Pteropus Alecto*. We used the complementary RNAi and CRISPR libraries to interrogate *Pteropus Alecto* cells for infection with two different viruses, mumps virus and Influenza A virus, respectively. Screening results converged on the endocytosis pathway and the protein secretory pathway as required for both viral infections. Additionally, we revealed a general dependence of the C-1-tetrahydrofolate synthase gene, MTHFD1, for viral replication in bat cells as well as in human cells. MTHFD1 inhibitor carolacton potently blocked replication of several RNA viruses including SARS-CoV-2. Our studies provide a resource for systematic inquiry into the genetic underpinnings of bat biology and a potential target for developing broad spectrum antiviral therapy.

---

As the only mammal capable of sustained flying, bats are intriguing animals with unique behavioral and physiological characteristics. Bats are notorious for transmission of deadly viruses, causing the epidemics of the Ebola^1^, SARS^2^, Henipavirus^3^, MERS^4^ and COVID-19^5,6^. Bats are widely considered the largest reservoir of numerous virus species but are resilient to the diseases associated with many of these viruses^7,8^. How bats tolerate the infections with these viruses is of intense interest due to the scientific and public health relevance. Genomic sequencing studies have revealed several genetic adaptions underlying this tolerance of viral infections. For example, the DNA damage checkpoint pathway and the NF-κB pathway are known to be positively selected in the bat genomes^9^. In addition, in the type I interferon gene family, the MHC class I genes and natural killer cell receptor family have been reported to be highly expanded in the bat genomes^10^. These studies provided evidence to support the hypothesis that bats have evolved a highly specific genetic configuration, especially in the innate immunity pathways. A major caveat is the lack of systematic examination of the functions of these genes due to shortage of functional genomic tools.

CRISPR and RNAi technologies have enabled genome-wide screening of gene functions for a variety of cell types, yielding an unprecedented wealth of information revealing the complexity of biological processes^11,12^. This is especially true in the field of virus-host interactions, where a plethora of host factors have been identified for a variety of viruses^13–15^. Both technologies have advantages and disadvantages and provide complementary approaches to achieve comprehensive genetic perturbations^16–18^. CRISPR permits complete knockout of a gene, thus affords generation of truly null genotype. A caveat is that the lethality associated with knockout of essential genes prevents functional parsing of these genes in different biological processes. In addition, CRISPR screening is usually used a pool-based format that requires the generation of stable cell lines through long term culturing. Therefore, CRISPR is suited for studying phenotypes that require a longer time to manifest. Based on previous studies, CRISPR screens demonstrated excellent specificity but limited sensitivity^17^. In contrast, siRNA screening relies on temporary knockdown of genes in an individual well format, a methodology suited for examining short-term phenotypic changes. Due to the transient nature of the assay, siRNA screening can tolerate growth defects due to gene knockdown and can elucidate hypomorphic phenotypes induced by siRNAs with varying degrees of knockdown. In general, RNAi screens allow comprehensive identification of genes in a biological process at the cost of more false positives due to pervasive off-target effects^17^. To fully realize the power of CRISPR and RNAi screening as complementary strategies to interrogate gene function in bats, here we developed a pool-based CRISPR library and a plate-based siRNA library for the whole genome of the model megabat *Pteropus Alecto.* We screened two viruses, influenza A virus and mumps virus, using the CRISPR and RNAi libraries, respectively, in the hope of uncovering common host factors for different viruses. From these two screens, common pathways such as the endocytosis pathway and protein secretory pathway have been identified, demonstrating conserved pathways between bat and other mammals in hosting viral infections. We extensively characterized a common hit from the two screens, MTHFD1, and identified it as a critical factor for viral replication in both bats and humans. We further demonstrate that MTHFD1 is a potential target for developing broad-spectrum antiviral drugs. MTHFD1 inhibitor carolacton demonstrated broad antiviral activities against Zika virus, mumps virus, and importantly, SARS-CoV-2. In summary, we developed two orthogonal libraries for genome-wide loss-of-function screening in bats and demonstrated their utility in discovery of important host factors for viral infections.

## Design and validation of genome-wide CRISPR of *P. Alecto*

We designed and synthesized the CRISPR and RNAi libraries based on the genome of *P. alecto.* For the CRISPR library, we adopted the algorithm developed for the construction of the popular GeCKO library to select single guide RNAs (sgRNAs)^19^. We designed 85,275 sgRNAs targeting 21,334 genes annotated in the *P. alecto* genome with 21,330 genes with 4 sgRNAs for each gene. The other 34 genes have 1-3 sgRNAs due to the constraints of the gene sequences for designing sgRNAs. Selection of sgRNAs was based on the calculations of off-target scores to lower potential off-target effects of the sgRNAs. The sgRNAs designs were validated for 21 sgRNAs targeting 11 genes using T7E1 endonuclease cleavage assay or Western blotting. We observed 90.5% of effectiveness in gene editing or gene knockdown, supporting the quality of the designed library (**Supplementary Figure 1**).

## CRISPR screening identified influenza A virus host factors

We performed a CRISPR screen with influenza A virus in PaKi cells to identify viral host factors (**Fig. 1a**). PaKi cells stably expressing SgCas9 and sgRNA library were infected with the H1N1 PR8 strain of influenza A virus, in duplicate. We used a high multiplicity-of-infection (MOI) so that naïve PaKi cells all died before the 9th day post infection. We then collected the surviving cells and compared sgRNA abundance with mock-infected cells. The two replicates demonstrated a high correlation, indicating high reproducibility (**Supplementary Fig. 2**). We calculated the results using the RIGER algorithm and identified 21 host factors required for the viral infection or pathogenesis^20^ (**Fig. 1b**). Three of the top four hits belong to the V-type proton ATPase subunits (ATP6V1B2, ATP6V1F and ATP6V0D1), which are involved in the acidification of endosome and have previously been previous identified as key host factors for influenza virus entry in several screens performed in human cells (**Fig. 1b**)^21^. Among the other hits were two genes, COPZ1 and SEC23B, involved in the protein secretory pathway, which are also known to be required for the replication of many viruses including influenza virus and flavivirus (**Fig. 1b**)^21–23^. The enrichment of previously known host factors in the identified hits in the CRISPR screen validated our library and the screening methodology.

**Figure 1.**
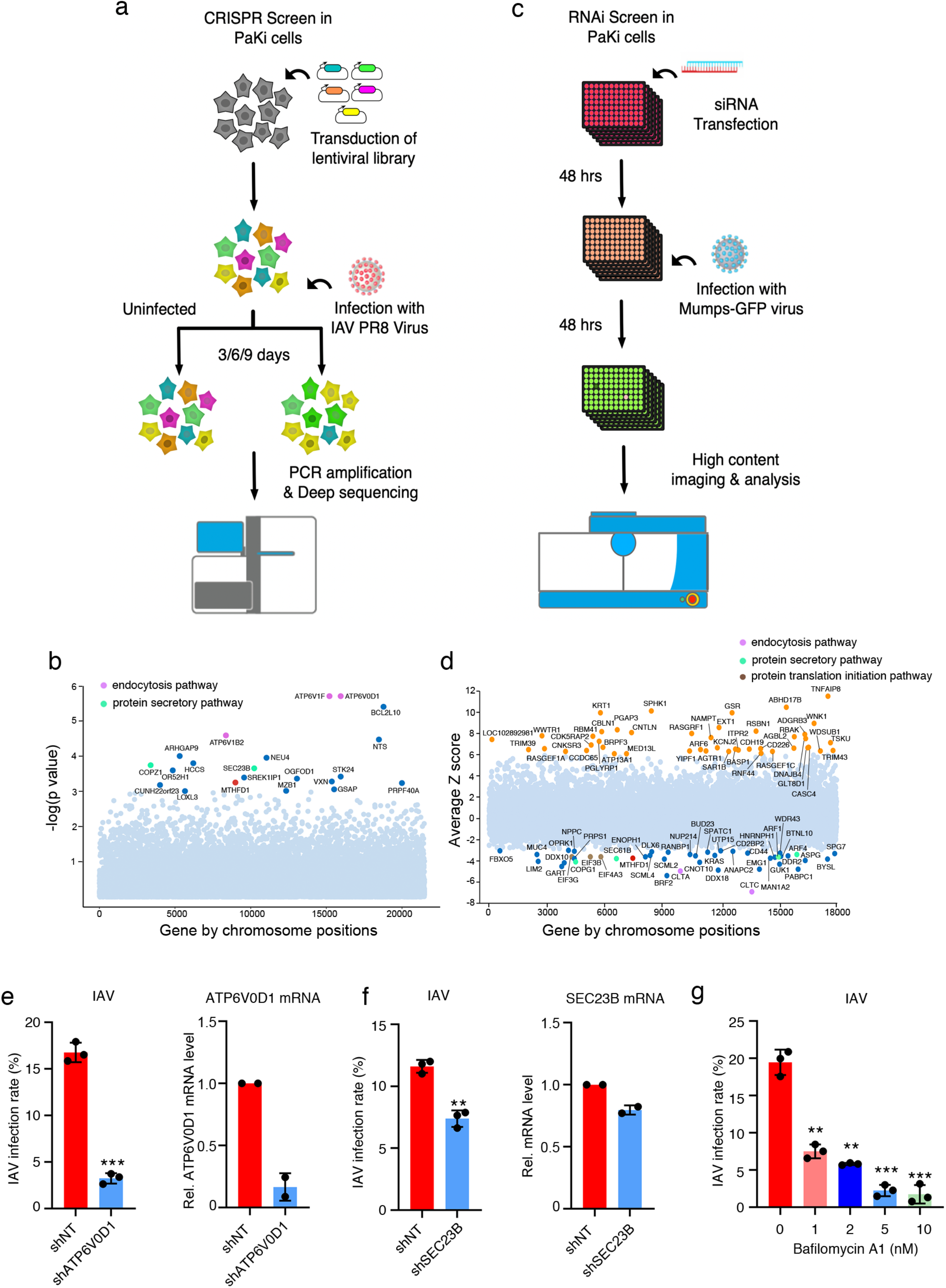
Work flows and results of CRISPR and RNAi screens in PaKi cells. **a**. For CRISPR Cas9 knock out screen, PaKi cells stably expressing Cas9 were transduced with lentiviral sgRNA library targeting 21300 genes of *P. alecto.* The transduced cells were selected with puromycin to generate the sgRNA cell library. Then the library cells were infected with influenza A virus (IAV) PR8 strain (A/PuertoRico/8/1934) at lethal dose or mock infected. The surviving cells were subjected to two more rounds of infection. The surviving cells at day 9 post-infection were harvested and the genomic DNA were prepared and amplified for the sgRNA region for deep sequencing. **b**. The sgRNA abundance from sequencing was analyzed by RIGER algorithm, the top hits representing host factors required for cell deaths from influenza infection were labeled. **c**. For siRNA screen, siRNAs targeting 18328 genes of *P. alecto* were arrayed in the 96-well plates. The siRNAs were reverse transfected into PaKi cells and 48 h after transfection, the cells were infected with mumps virus. The percentage of infected cells was calculated at 48 h post infection using high content imaging. **d.** The siRNA targeted genes were ranked by robust Z score, and the potential host factors or restriction factors were shown in the bottom or the top, respectively. **e**. PaKi cells expressing an shRNA targeting ATP6V0D1 or a non-targeting control shRNA were infected with IAV and the virus infection rate was measured by immunofluorescence using the hemagglutinin antibody at 12h post infection (left). The shRNA knockdown efficiency was detected by real-time PCR (right). For immunofluorescence experiments, n=3, and for real-time PCR data, n=2. Data shown as mean ± standard deviation. f. PaKi cells expressing an shRNA targeting SEC23B or a non-targeting control shRNA were infected with IAV PR8 and the virus infection rate was measured by immunofluorescence using the hemagglutinin antibody at 12h post infection (left). The shRNA knockdown efficiency was detected by real-time PCR (right). For immunofluorescence experiments, n=3, and for real-time PCR data, n=2. Data shown as mean ± standard deviation. **g.** PaKi cells were treated with indicated doses of V-ATPase inhibitor, bafilomycin A1, and infected with IAV. The infection rate was measured by immunofluorescence using the hemagglutinin antibody at 12h post infection. Data shown as the mean ± standard deviation (n=3). *: P<0.05, **: P<0.01, ***: P<0.001.

## RNAi screen results converge with CRISPR screening on key pathways

We synthesized the sgRNA library in an array-based oligonucleotide library and cloned them into the lentiGuide-Puro lentivirus expression vector^24^. A custom bat siRNA library was designed to target 18,328 genes using 40,016 siRNA duplexes. We performed RNAi screening in PaKi cells for identification of host factors for the infection of mumps virus, a paramyxovirus (**Fig. 1c**). The screen was performed in 384-well format with a mumps virus strain expressing EGFP from an additional open reading frame^25^. PaKi cells were infected 48 hours after transfection of the siRNA library. High-content imaging analysis was utilized to quantitate the infection rate as well as the total cell number each well, indicating cytotoxicity of the siRNA. Excluding siRNAs with significant cytotoxicity, we selected 45 host dependency factors whose knockdown resulted in significantly reduced viral infection and 45 host restriction factors whose knockdown promoted viral infection based on the Z score of infection rate (**Fig. 1d**). We selected a number of genes for individual validation with more replicates and obtained a high validation rate (11/12 of host dependency factors and 7/12 of the host restriction factors validated with at least one siRNA) (**Supplementary Fig. 3**), supporting the reproducibility of our screening methodology.

Although there is only one gene overlap between the host factors identified in both CRISPR and RNAi screens, we observed that the endocytosis pathway and protein secretory pathway were enriched in both screens, albeit represented by different genes (**Fig. 1b and 1d**). For the endocytosis pathway, two genes encoding components of clathrin complex, CLTA and CLTC, were among the top three hits (**Fig. 1d**). Four genes involved in the protein secretory, namely ARF1, ARF4, COPG1 and SEC61B, were identified (**Fig. 1d**). This is consistent with the idea that both influenza A virus and mumps require the two pathways for viral replication. In addition, protein translation initiation, which was previously reported to be required for influenza A infection^21^, was also represented in the top hits by three genes: EIF3B, EIF3G and EIF4A3. Conversely, RNAi screening revealed genes that restrict mumps infection in PaKi cells. Among the top hits were several known antiviral genes, TRIM39, TRIM43^26^ and NAMPT^27^. Overall, we identified many candidate genes that might play a role in viral infection in bats.

The two pathways identified by both CRISPR screen and siRNA screen, endocytosis and protein secretion, have previously been implicated in human host response to a number of viral^13,21,23^ pathogens To validate the role of these pathways in the bat response to viral infection, we established PaKi stable cell lines expressing shRNAs targeting several hits from the two screens. Stable knockdown of a V-type proton ATPase subunit and SEC23B led to a significant decrease of influenza A virus infection in PaKi cells (**Fig. 1e and 1f**). In addition, the small molecule inhibitor of V-type proton ATPase, bafilomycin A (**Fig. 1g**), potently blocked influenza infection. Taken together, these results demonstrate that bat and human cells share key genes and pathways in hosting viral infection.

## MTHFD1 is a key host factor for a broad spectrum of RNA viruses

The gene identified as a host factor from both the CRISPR and RNAi screen was MTHFD1 (**Fig. 1c and 1d**), which we validated by siRNA knockdown in the context of mumps infection (**Supplemental Fig. 3**). We validated its function in influenza A virus replication in PaKi cells with two different sgRNAs targeting MTHFD1 (**Fig. 2a–2b**). Similarly, in HEK293T cells with MTHFD1 knocked out by two sgRNAs, influenza replication is also inhibited, suggesting a conserved requirement for this gene by influenza virus in human cells (**Fig. 2c**). Further virus titration assay demonstrated that MTHFD1 knockdown significantly inhibits viral replication for mumps virus in PaKi cells (**Fig. 2d and 2e**). The effect is similar to the knockdown of SEC61B, a well-studied host protein required for viral replication and a top hit in our RNAi screen (**Fig. 2d and 2e**)^23^. We additionally tested Melaka virus (PRV3M), a pteropine orthoreovirus that causes acute respiratory disease when transmitted by bats to humans^28^. PRV3M titer was also significantly reduced by MTHFD1 knockdown (**Fig. 2e and 2f**). Infection of Zika virus, a member of the flavivirus family, was also inhibited by MTHFD1 knockdown using siRNA (**Fig. 2g**). Alternative knockdown with stable expression of two shRNAs validated the requirement of MTHFD1 for the replication of mumps virus and Zika virus (**Fig. 2h, 2i and 2j**). Importantly, we also show that this inhibition is completely rescued by overexpression of MTHFD1 from a construct resistant to the shRNA, ruling out potential off-target effects (**Fig. 2k**).

**Figure 2.**
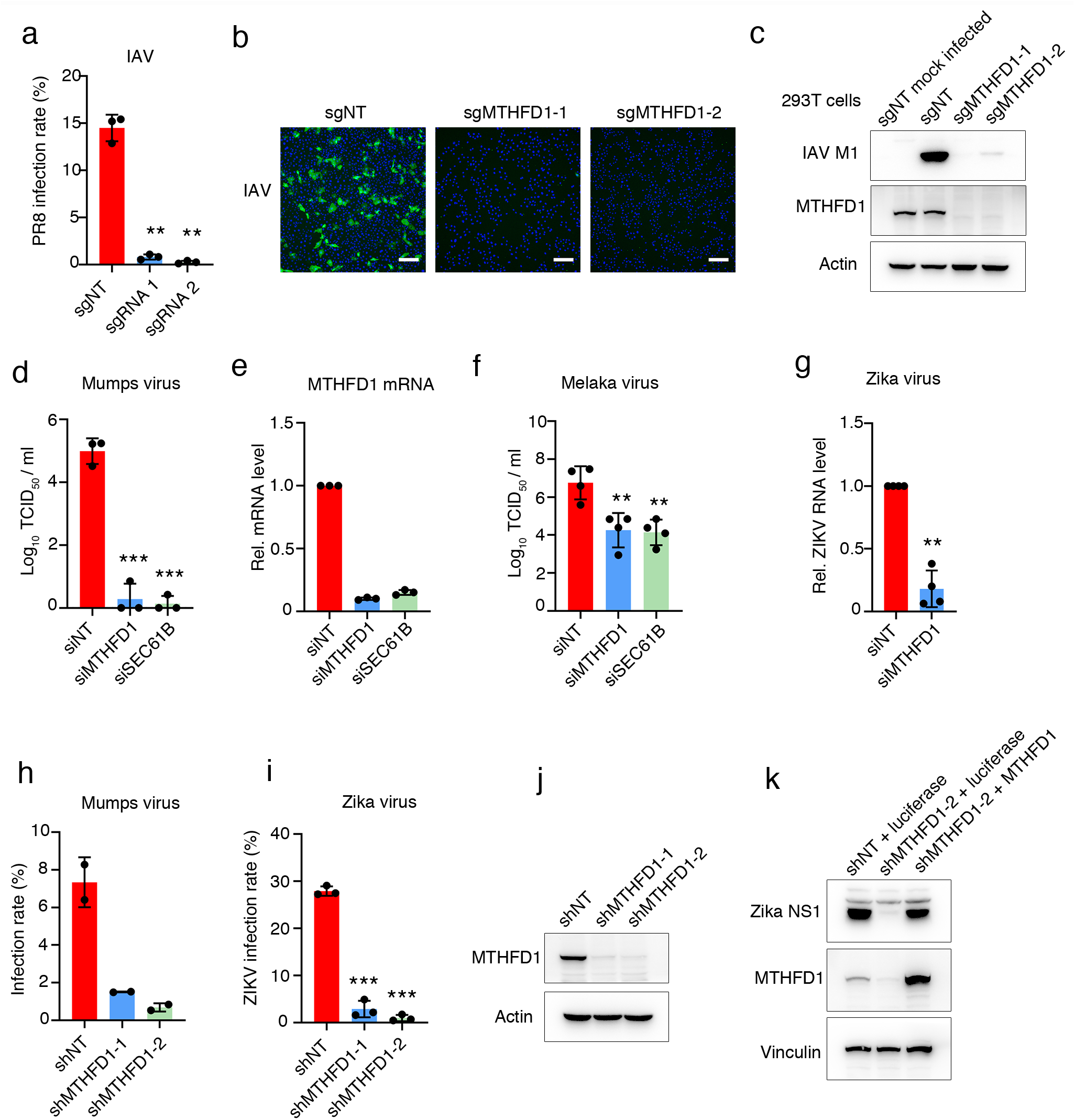
MTHFD1 is host factor for replication of multiple RNA viruses. **a.** MTHFD1 knockout clonal PaKi cells were generated from two independent sgRNAs and infected with Influenza A virus (IAV) for 12h. The infection rates were quantified with immunofluorescence staining of HA. Data shown as the mean ± standard deviation (n=3). **b.** Representative images from **a.** (scale bar=100 μm). **c**. MTHFD1 knock out 293T cells were generated from two independent sgRNAs and infected with IAV for another 12h. Then the cells were collected and analyzed by Western blotting. Data shown are representative of three independent experiments. **d-f.** PaKi cells were transfected with non-target control siRNA (siNT) or siRNAs targeting MTHFD1 and SEC61B followed by mumps virus(**d**) and Melaka virus **(f)** infection. The virus in the supernatant were tittered by limiting dilution. The knockdown efficiency was determined by realtime PCR and shown in **(e).** For **d** and **e** n=3, and for **f**, n=4. Data shown as the mean ± standard deviation. **g**. PaKi cells were transfected with siNT or siRNA targeting MTHFD1, 48 h later the cells were infected with Zika virus. The Zika virus RNA level was measured by real-time PCR. For **g,** n=4. Data shown as the mean ± standard deviation. **h-j**. PaKi shRNA control cells (shNT) and shMTHFD1 cells were infected with mumps virus **(h)** or Zika virus **(i)**, the virus infection level was measured by immunofluorescence. For **h**, n=2 and for **i**, n=3. Data shown as the mean ± standard deviation. The shRNA knockdown efficiency was tested by Western blotting as showing in (**j**). Data shown are representative of three independent experiments. **k**. Luciferase or shRNA resistant MTHFD1 construct were stably expressed in shMTHFD1-2 PaKi cells, then the cells were infected with Zika virus. The rescue level of Zika virus was analyzed by Western blotting. Data shown are representative of three independent experiments. For all bar graphs, the mean ± standard deviation values are shown. *: P<0.05, **: P<0.01, ***: P<0.001.

## MTHFD1’s formyl tetrahydrofolate synthetase activity is essential for RNA virus replication

MTHFD1 is a trifunctional enzyme involved in the one carbon (C1) metabolism pathway, which is responsible for cellular production of purine, dTMP and methyl groups^29^. MTHFD1 has three enzymatic functions including dehydrogenase and cyclohydrolase activities encoded in the N-terminal domain and formyl tetrahydrofolate (Formyl THF) synthetase activity encoded in the C-terminal domain (**Fig. 3a and 3b**). We suspected that MTHFD1 knockdown would lead to the deficiency of cellular purine levels thereby reducing RNA replication. Indeed, supplementation of purine analogs, either hypoxanthine or inosine in the media almost completely rescued the replication of influenza A and Zika viruses in MTHFD1-knockout or knockdown PaKi cells as shown by both immunofluorescence staining and virus tittering assay (**Fig. 3c, 3d and 3e**). In contrast, supplementation of methyl-group donor or thymidine had little effect on viral replication, evidence that it is the purine synthesis activity of MTHFD1 that is essential for the viral replication (**Fig. 3d**). To further pinpoint the enzymatic function of MTHFD1 in viral replication, we performed gene complementation experiments in an shRNA cell line with over 90% knockdown of MTHFD1 (**Fig. 2i**). The block of Zika virus replication in this cell line was completely rescued by overexpression of an shRNA-resistant MTHFD1 construct or bat homolog (**Fig. 3f and 3g**). Mutations of MTHFD1 that are known to disrupt its dehydrogenase and cyclohydrolase activities (D125A and R173C)^30,31^ did not affect this rescue (**Fig. 3b, 3f and 3g**). However, mutations in the Formyl THF synthetase active site (F759A/W788A, or FW/AA)^32^ completely abolished this rescue, further supporting that the purine synthesis activity of MTHFD1 is an essential activity for viral replication (**Fig. 3b, 3f and 3g**).

**Figure 3.**
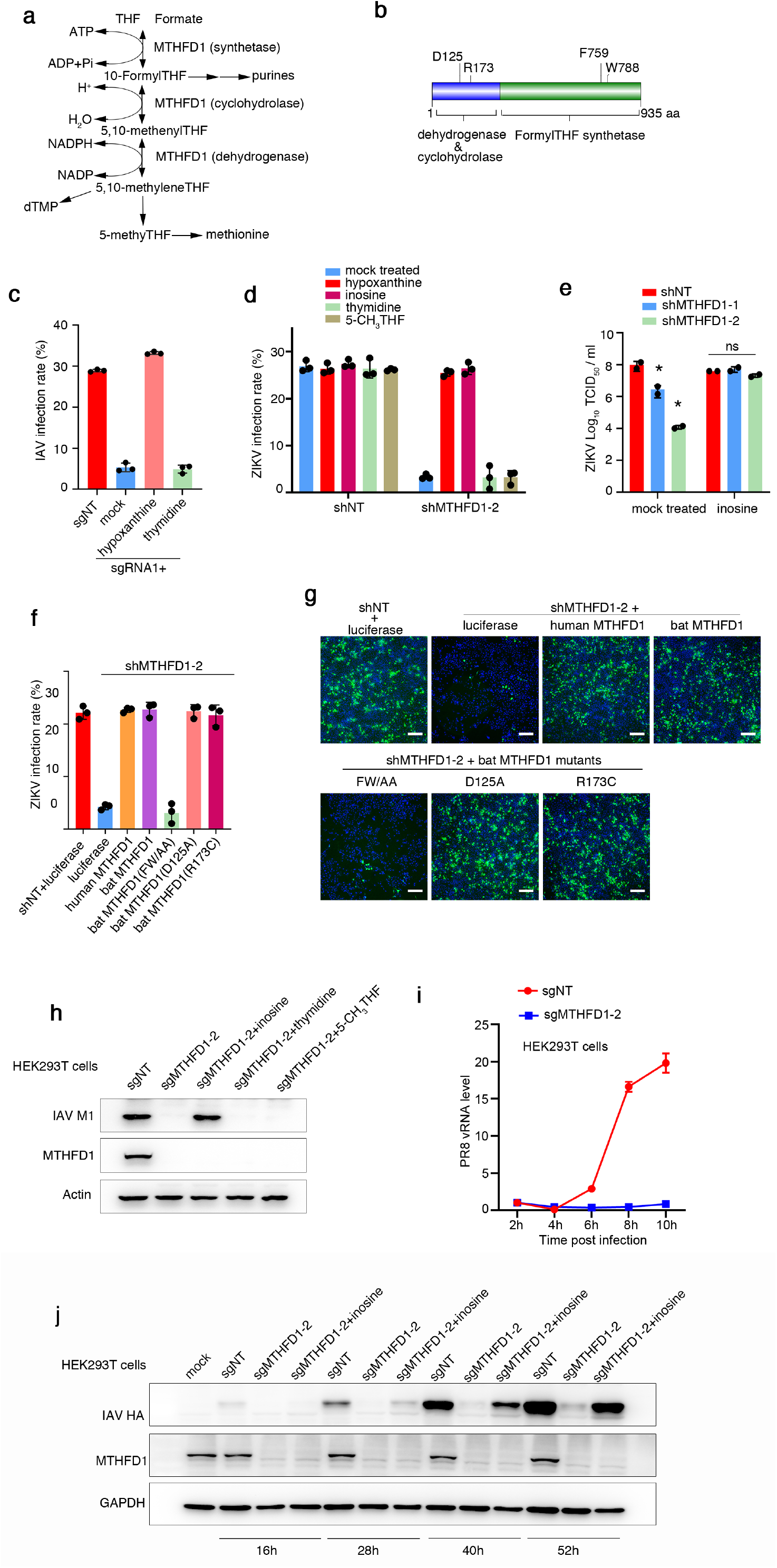
MTHFD1’s formyl tetrahydrofolate synthetase activity is essential for RNA virus replication, **a.** MTHFD1 is a trifunctional enzyme in the one carbon metabolism which is required for purines, dTMP and methionine synthesis. **b**. Domain structures and key amino acids of MTHFD1 with key residues for the three enzymatic functions indicated. **c**. PaKi MTHFD1 knock out cells were infected with IAV with or without supplementation of hypoxanthine or thymidine for 12 h and the virus infection level was detected by immunofluorescence. Data shown as the mean ± standard deviation (n=3). **d**. PaKi shMTHFD1-2 cells or control cells with a non-targeting shRNA were infected with Zika virus in regular medium or in medium supplemented with inosine, thymidine or 5-CH_3_THF. Data shown as the mean ± standard deviation (n=3). **e**. PaKi shMTHFD1 cells were infected with Zika virus in regular medium or in medium supplemented with inosine. At 48 h post infection, the virus in the supernatant was harvested and the titre was determined by limiting dilution in PaKi cells. Data shown as the mean ± standard deviation (n=2). f. PaKi shMTHFD1-2 cells or control cells with a non-targeting shRNA were transfected with luciferase (negative control), human wild type MTHFD1, bat wildtype MTHFD1 or bat MTHFD mutants, followed by Zika virus infection. The virus infection rate was measured by immunofluorescence. Data shown as the mean ± standard deviation (n=3). **g.** Representative images from (**f**). Scale bar: 100 μm. **h.** HEK293T MTHFD1 knock out cells or control HEK293T cells with a non-targeting sgRNA (sgNT) were infected with PR8 with or without the supplementation of inosine, thymidine or 5-CH3THF. At 12 h post infection, the cells were collected and analyzed by Western blotting. Data shown are representative of three independent experiments. **i**. HEK293T MTHFD1 knockout cells (sgMTHFD1-2) or control cells (sgNT) were infected with IAV. The vRNA level was determined by real-time PCR at the indicated time points. The results shown representative of three independent experiments **j.** HEK293T MTHFD1 knockout cells (sgMTHFD1-2) or control cells (sgNT) were transfected with plasmids expressing the influenza virus mini-genome including polymerase subunits PB1, PB2 and PA, and nucleoprotein together with pPOLI-HA, which can transcribe the hemagglutinin vRNA of influenza virus with or without inosine supplementation. At 16h, 28h, 40h and 52h post transfection, the cells were collected and analyzed by Western blotting. Data shown are representative of three independent experiments. For all bar graphs, the mean ± standard deviation values are shown. *P<0.05.

We observed a similar dependence on MTHFD1 function for influenza A virus infection in HEK293T cells, a human kidney epithelial cell line (**Fig. 3h**). Time course experiments demonstrated a viral replication following MTHFD1 knockdown steadily decreased from four hours post-infection with influenza A virus, consistent with a RNA replication defect (**Fig. 3i**). This defect in replication was completely rescued by supplementing purine analog inosine in the media, supporting a conserved function of MTHFD1 in viral infection in bats and humans (**Fig. 3h and 3j, Supplementary Fig. 4**).

## MTHFD1 inhibitor carolacton inhibits RNA virus replication

Carolacton, a natural product derived from bacteria, has recently been identified as a potent inhibitor of MTHFD1^33^. We tested the effect of this compound on viral replication. Carolacton inhibited Zika virus and mumps virus replication in PaKi cells in a dose dependent fashion (**Fig. 4a, 4b and 4c**). The inhibition was achieved at doses with little cytotoxicity (**Fig. 4a**). Mechanistically, this inhibition was completely rescued by supplementation of inosine, providing support that carolacton indeed exerts antiviral activities by inhibiting MTHFD1 (**Fig. 4b and 4c**). We next tested the effect of carolacton on the infection of SARS-CoV-2. The viral infection in Vero E6 cells was effectively inhibited at sub-micromolar concentrations by carolacton (IC_50_=0.14 μM) (**Fig. 4d**). The cytotoxic effect is very moderate in comparison (**Fig. 4d**), supporting a therapeutic window of this compound in potential clinical applications.

**Figure 4.**
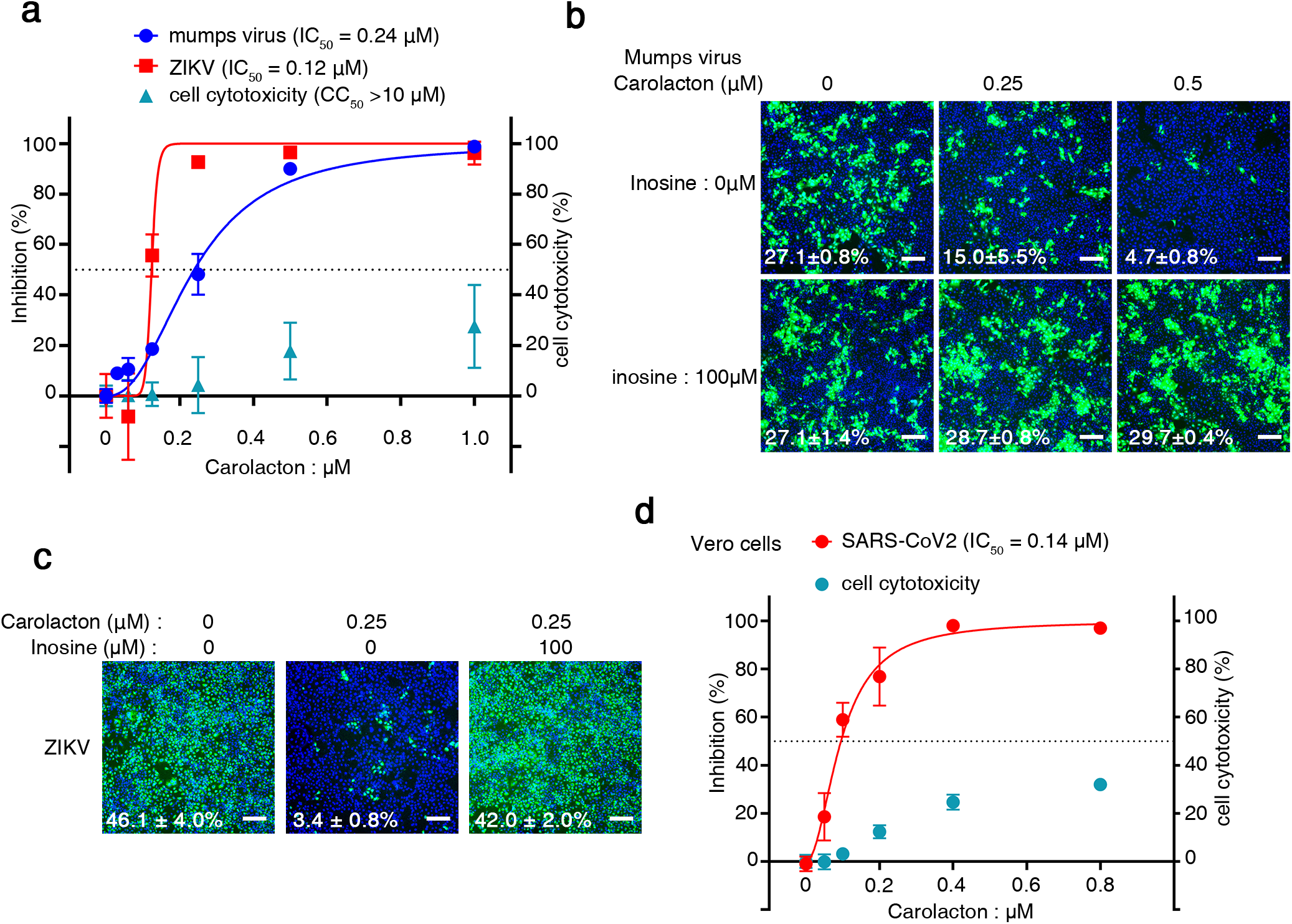
MTHFD1 inhibitor carolacton can inhibit a number of RNA viruses including SARS-CoV-2. **a.** PaKi cells were treated with different concentrations of carolacton and infected with mumps virus expressing EGFP or Zika virus. Virus infection level was detected by EGFP or immunofluorescence staining of Zika E protein. The cell viability was measured by MTT assay. n=3. Data shown as the mean ± standard deviation. **b.** Representative images of immunofluorescence staining of mumps infection from (**a**). **c.** PaKi cells were treated with carolacton and infected with Zika virus in the presence of inosine or mock treated. Representative images of immunofluorescence staining of Zika virus E protein were shown. Infection rate shown as the mean ± standard deviation (n=3). **d.** PaKi cells were treated with carolacton, and 1h later infected with SARS-CoV-2. At 48 h post infection, the SARS-CoV-2 in the supernatant was measured by real-time PCR. Data shown as the mean ± standard deviation (n=3).

## Discussion

We developed the first bat genome-wide RNAi and CRISPR libraries and performed two screens with two different RNA viruses. Although the screening methodologies are quite different, we uncovered similar pathways essential for viral replication, namely the endocytosis pathway and protein secretory pathways. The different genes in these two pathways identified from the screens reflect the complementarity of the methodologies, with RNAi screening useful for observing acute effects on viral infection and CRISPR screening for long-term effects on cell survival. Other than technical differences between screening methods, viral kinetics, sensitivity of the viruses to loss-of-function of genes, differences in genome coverage within the two libraries likely contributed to the different hits observed in our studies. Both libraries provide valuable and complementary sources that meet the urgent need to address the biological questions of bats. Similar to human genomic libraries, the utility of these bat libraries in screening different cell types of bats, especially those of the immune systems, would unveil more secrets of this enigmatic species.

Both screens identified MTHFD1 as an essential viral host factor. MTHFD1 is a key enzyme in the C1 metabolism responsible for production of purine, dTMP and methyl group, the first two of which are building blocks for DNA and RNA synthesis. Interestingly, all previous RNAi and CRISPR screens using human cells have not identified MTHFD1 as a host factor. Moreover, we found bats cells and tissues have lower MTHFD1 expression levels than their human counterparts (**Supplementary Fig. 5a, 5b and 5c**). This might be related to the difference in the promoter regions between bats and humans, notably a lack of CCAAT box^34^ in the promoter of two bat species (**Supplementary Fig. 5d).** We show that MTHFD1 knockdown blocks replication of multiple RNA viruses, including influenza virus, mumps virus, Melaka virus and Zika virus. Importantly, host cells have a higher tolerance for MTHFD1 inhibition than viruses, potentially providing a therapeutic window for targeting MTHFD1 with antiviral drugs. Virus infection converts the whole cellular machinery into a factory to massively produce progeny viruses, which explains the vital dependence on MTHFD1. This is analogous to cancer cell’s addiction to non-oncogenic gene functions such as many metabolic enzymes^35^. Similar to the therapeutic targeting of these non-oncogenic addictions for developing cancer drugs, these viral addictions to metabolic enzymes such as MTHFD1 could open doors to many new opportunities for developing broadspectrum antiviral drugs^36^. These drugs could be readily applied in emerging epidemics, as demonstrated by our evidence here that MTHFD1 inhibitor carolacton can suppress the infection of SARS-CoV-2. We envision the functional genomics tools we provided here will be instrumental to further discoveries by better understanding the bat-virus interaction.

## Author contributions

XT, DEA and L-FW conceived and guided the study. DEA performed and analyzed RNAi screen and RNAi hit validation with assistance from SYK, BGY, KS and XFL. JC and QY performed CRISPR screen and hits validation. JC performed most experiments on MTHFD1. BH performed SARS-CoV-2 infection assay with help from FY and PFY and under the guidance of WT. WZ, JG, WL and XZ performed bioinformatics analysis of the study. XT wrote the manuscript with input from all authors.

## Acknowledgements

We thank Fu Chengzhang and Rolf Müller for providing carolacton. We thank Jianzhong Xi for oligonucleotide synthesis. XT is supported by the the China National Funds for Excellent Young Scientists (31722030) and was supported by the Beijing Advanced Innovation Center for Structural Biology and the Tsinghua-Peking Joint Center for Life Sciences. DEA and L-FW are supported by grants NRF2012NRF-CRP001–056 and NRF2016NRF-NSFC002-013 from the Singapore National Research Foundation and National Medical Research Council Cooperative Basic Research Grant NMRC/BNIG/2030/2015.

## Methods

### Cells and virus

PaKi, Vero and 293T cells were cultured in DMEM supplemented with 10% heat inactivated FBS. Influenza H1N1 PR8 (A/Puerto Rico/8/1934) was propagated in embryonated eggs. Mumps virus was propagated in PaKi or Vero cells.

### Genome-wide CRISPR-Cas9 knockout screen

Cas9 stably expressed bat kidney derived PaKi cells were transduced with lentivirus of sgRNA library targeting 21300 genes in the genome of *P. alecto.* The transduced cells were selected with puromycin to generate the library cells. Then the sgRNA library cells were mock infected (negative control) or changed with influenza A H1N1 PR8 strain (A/Puerto Rico/8/1934) at lethal dose. The surviving cells were subjected to two more round infection. The final surviving cells were applying to deep sequencing. And the sgRNA abundance was analyzed by RIGER algorithm^1^.

### Deep sequencing data analysis

De-multiplexed reads were trimmed by Cutadapt^2^ (version 1.18) using 22 bp flanking sequences around the guide sequence. Trimmed reads were mapped to the index of bat sgRNA guide library using Bowtie2^3^. Samtools^4^ (version 1.9) was used to process the alignment files to retain perfectly mapped reads only, and a custom-made Python script was used to count the reads mapped to each guide. The reads count of guide for each condition were then normalized using DEseq2. The log2 ratio of counts between conditions were calculated. The log2 fold change of reads count of infected samples to mock infected samples were then used as the input for RIGER algorithm as previously described (RNAi Gene Enrichment Ranking, GENE-E)^1^. The log-fold-change metric for ranking sgRNAs and the weighted sum method (weighted sum of first two ranks of sgRNAs for a gene) with 1 × 10^6^ permutations were used to convert individual sgRNA to genes.

In the output of RIGER analysis, a given gene was assigned a normalized enrichment score (NES) to give the ranked gene list. Hits were defined as those owning p-value smaller than 0.001.

### siRNA screen

For siRNA screen, siRNAs targeting 18,328 genes of *P. alecto* were arrayed in the 96-well plates. The siRNAs were reverse transfected into PaKi cells, 48 h later, the cells were infected with mumps virus. The virus infection level was measured at 48 h post infection by high content imaging (Operetta) as previously described^5^. The siRNA targeted genes were ranked by robust Z scores^6^, and the potential host factors (Z score < −3) or restriction factors (Z score > 6) were identified.

### Validation of sgRNA

Single sgRNA was cloned into lentiGuide-puro and packaged in lentivirus, which was transduced into Cas9 stably expressing PaKi cells, followed by puromycin and blasticidin selection. The clonal cells were generated by limiting dilution. The knock out or knock down efficiency was tested by Western blotting. To facilitate the growth of single clone MTHFD1 knock out cells, the cells were cultured with HT medium (DMEM supplemented with 10% FBS, 100μM hypoxanthine and 16μM thymidine).The knock out or knock down efficiency was tested by Western blotting. Validated knockout cells were applied to virus infection assays. For MTHFD1 knock out cells, the HT was removed 8h before virus infection.

### Validation of shRNA

Individual shRNAs were designed using the BLOC-iT^TM^ RNAi Designer (Thermo Fisher Scientific). The specificity was tested by BLAST (NCBI). The shRNA primes were synthesized as oligos and cloned into PLKO.1.

PaKi shMTHFD1-1 forward primer: 5’-ccgg GCACATGGGAATTCCTCTACCctcgagGGTAGAGGAATTCCCATGTGC tttttg-3’ PaKi shMTHFD1-1 reverse primer: 5’-AATTCAAAAA GCACATGGGAATTCCTCTACC CTCGAG GGTAGAGGAATTCCCATGTGC-3’ PaKi shMTHFD1-2 forward primer: 5’-ccgg GCCTGCTGTCACTTAGGAAATctcgagATTTCCTAAGTGACAGCAGGC tttttg −3’ PaKi shMTHFD1-2 reverse primer: 5’-AATTCAAAAA GCCTGCTGTCACTTAGGAAAT CTCGAG ATTTCCTAAGTGACAGCAGGC −3’

### Western blot

Cells were harvested and lysed in RIPA lysis buffer (Beyotime, P0013C) with a cocktail of protease inhibitors (Biomake, B14001). Samples were applied to 10% or 12% SDS/PAGE gels(BioRad) and transferred to PVDF membranes (millipore). Membranes were blocked with 5% nonfat milk in PBS with 0.1% tween 20. Then probed with primary antibodies. The following antibodies were used in this study: anti-MTHFD1 (Proteintech, 10794-1-AP), anti-beta actin (Easybio, BE0022), anti PR8 M1 (Genetex, GTX125928-S), anti HA (H1N1) (Genetex, GTX117951-S), anti-ZIKV NS1 (Genetex, GTX133307), anti flavivirus group antigen antibody (Millipore, MAB10216). The blots were developed by HRP reaction and imaged with a ProteinSimple FluorChem imaging system.

### RNA isolation and real-time PCR

Total RNA from the cells were isolated with a HiPure Total RNA Mini Kit (Magen, R4111-03). The cDNA was generated by RevertAid RT Reverse Transcription Kit (thermos scientific, K1691). Influenza vRNA specific primer was used for vRNA transcription. The quantitative realtime PCR was performed with SYBR Green qPCR master mix (Vazyme, Q311-02) and data were analyzed by Bio-Rad CFX96 system. The primers for target genes were as follows: Human GAPDH forward primer: 5’-ACAACTTTGGTATCGTGGAAGG-3’ Human GAPDH reverse primer: 5’ – GCCATCACGCCACAGTTTC-3’ P. alecto ACTIN forward primer: 5’ – gccagtctacaccgtctgcag −3’ P. alecto ACTIN reverse primer: 5’ – cgtaggaatccttctggcccatg-3’ P. alecto MTHFD1 forward primer: 5’- gggagcgactgaagaaccaag-3’ P. alecto MTHFD1 reverse primer: 5’- tcttcagcagccttcagcttcac-3’ P. alecto SEC23B forward primer: 5’- cagcgtttgaccaggaggcc-3’ P. alecto SEC23B reverse primer: 5’- gggtcagatcctgtcgggc −3’ P. alecto GAPDH forward primer: 5’- ATACTTCTCATGGTTCACAC −3’ P. alecto GAPDH reverse primer: 5’ – TCATTGACCTCAACTACATG-3’ P. alecto ATP6V0D1 forward primer: 5’ – GTGGTAGAGTTCCGCCACAT-3’ P. alecto ATP6V0D1 reverse primer: 5’ – CTCAAAGCTGCCTAGTGGGT-3’ PR8 M1 forward primer: 5’ – TTCTAACCGAGGTCGAAACGTACG-3’ PR8 M1 reserve primer: 5’- ACAAAGCGTCTACGCTGCAG-3’ PR8 vRNA reserve transcription primer: 5’-AGCRAAAGCAGG-3’ ZIKV NS5 forward primer: 5’- GGTCAGCGTCCTCTCTAATAAACG-3’ ZIKV NS5 reserve primer: 5’- GCACCCTAGTGTCCACTTTTTCC-3’

### Immunofluorescence

Virus infected cells were fixed with 4% paraformaldehyde (PFA) for 10min at room temperature (RT), and permeated with 0.2% Triton X-100 for another 10min at RT, the cells were washed with PBS for 3 times and incubated with PR8 HA antibody or ZIKV E protein antibody for 2h at RT or 4 °C overnight, then washed with PBS for 3 times and incubated with second antibody AF488 (goat anti mouse). The nuclei were stained with DAPI. Images were acquired with Cellomic ArrayScan VTI HCS (Thermo Scientific)

### Virus titration

Approximately 1×10^4^ PaKi cells were plated into each well of 96-well plates. The supernatant containing virus harvested from previously infected cell was subjected to ten-fold serial dilution. Then dilutions were added to the cells and cytopathic effect was observed 96 h after infection. Titres were determined using standard methodology^7^.

### Influenza virus mini-genome assay

Approximately 2×10^5^ of 293T control cells or 293T MTHFD1 knock out cells were plated into each well of 12-well plates and the knockout cells were cultured with HT (100 μM hypoxanthine and 16 μM thymidine, Gibco) medium. Then the cells were transfected with the plasmids expressing viral polymerase subunits PB1, PB2 and PA, and nucleoprotein, together with pPOLI–HA, transcribing hemagglutinin vRNA segment of A/WSN/33(H1N1) (WSN). At the time of transfection, HT was removed from the cells and supplied with fresh medium with or without inosine (100 μM). The HA protein level were analyzed at 16 h, 28 h, 40 h and 52 h post transfection by Western blot.

### MTT assay

MTT cell viability assay (Solarbio) was used to analyze cell viability according to the manufacturer’s instructions. Briefly, 10 μl MTT (0.5%) mixed with 90 μl fresh medium was added into the well. The cells were incubated at 37°C for 4 h, after the formazan crystals were formed, the medium was removed and 110 μl DMSO was used to solubilize the formazan crystals. The absorbance at 490 nm was measured with a multimode plate reader victory x5 (Perkin Elmer).

---

**Supplementary Figure 1.**
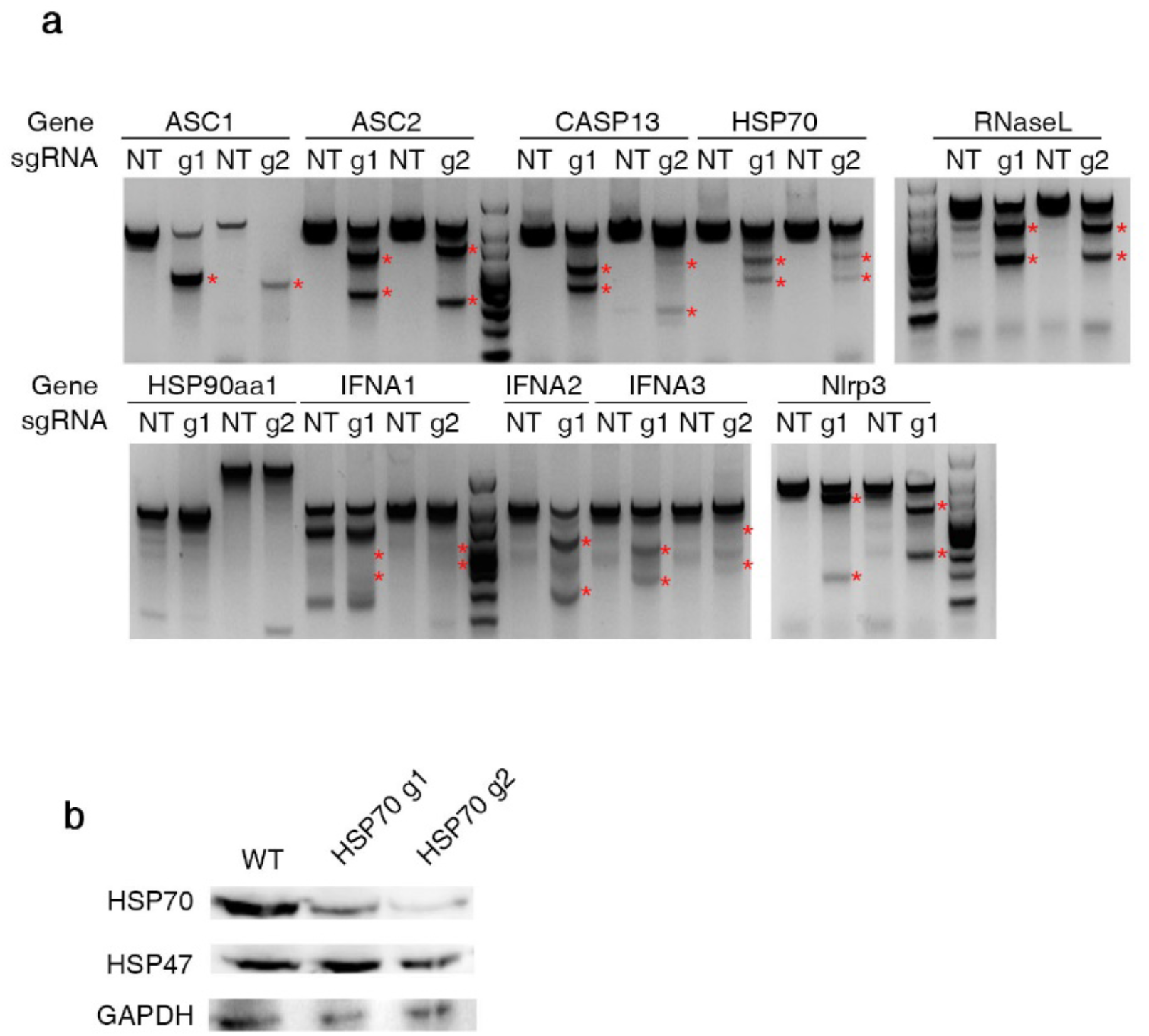
Validation of sgRNA design of ten genes in PaKi cell line using CRISPR-Cas9. **a.** T7E1 assay to confirm indels in the target genes. NT: nontargeting control, g1,g2: designed guide RNA targeting each gene. **b.** Western blot to confirm the decrease of protein level of HSP70 by two guide RNAs g1 and g2.

**Supplementary Figure 2.**
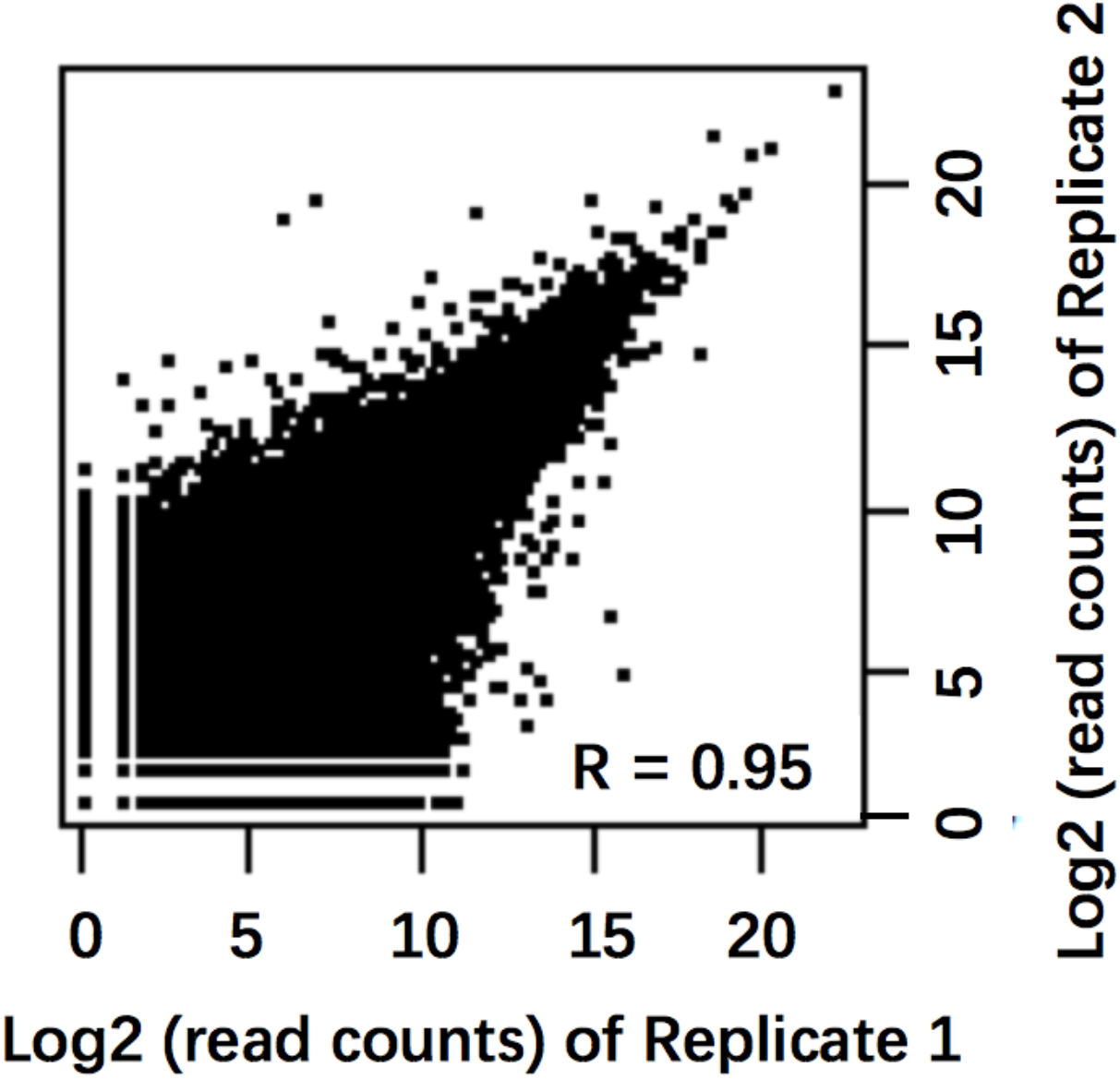
Correlation between the read counts of each sgRNA in two biological replicates of the CRISPR screen.

**Supplementary Figure 3.**
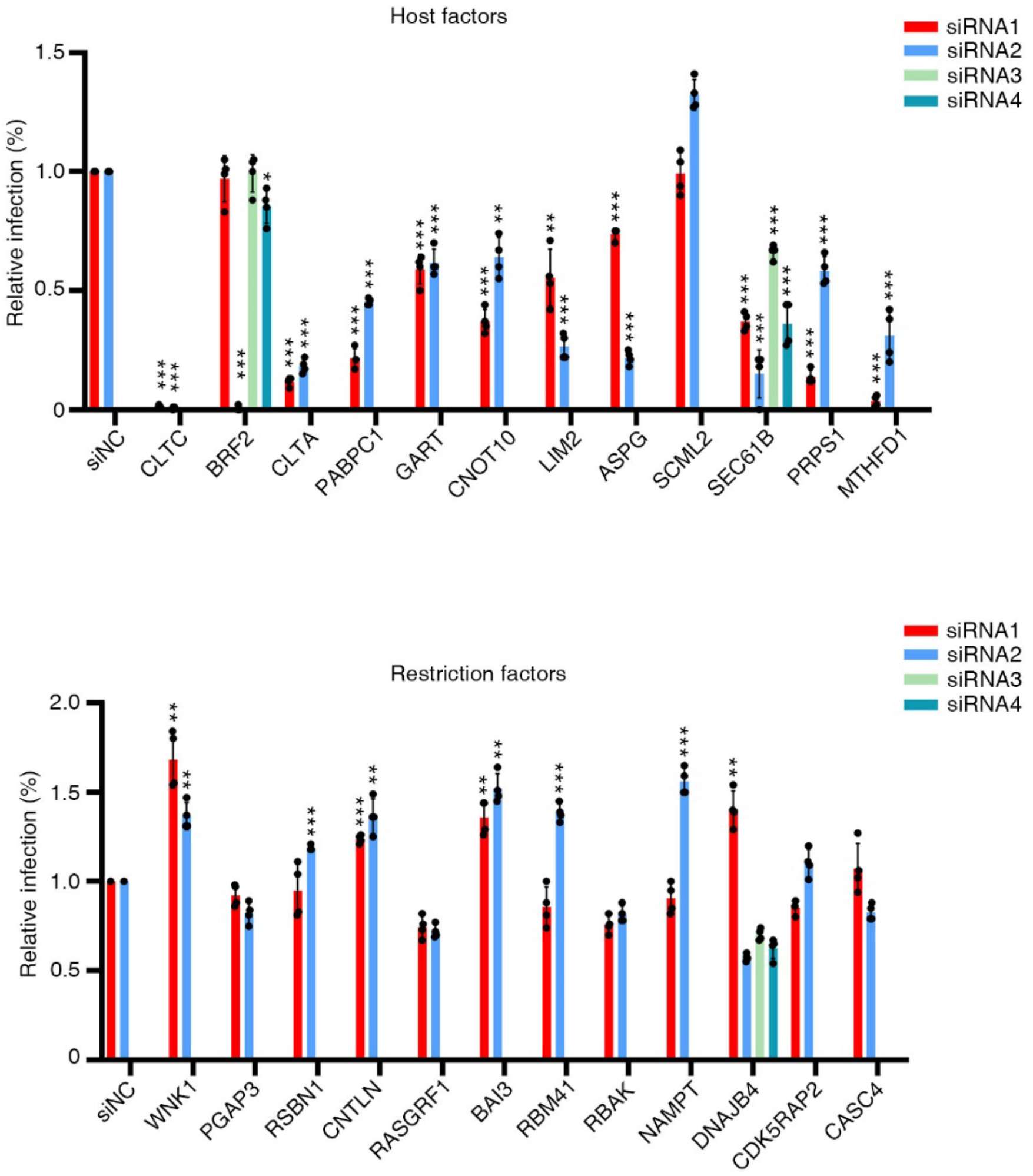
Validation of siRNA screen hits. PaKi cells were transfected with the siRNAs followed by mumps-GFP virus infection. The virus infection levels were measured by high content imaging.

**Supplementary Figure 4.**
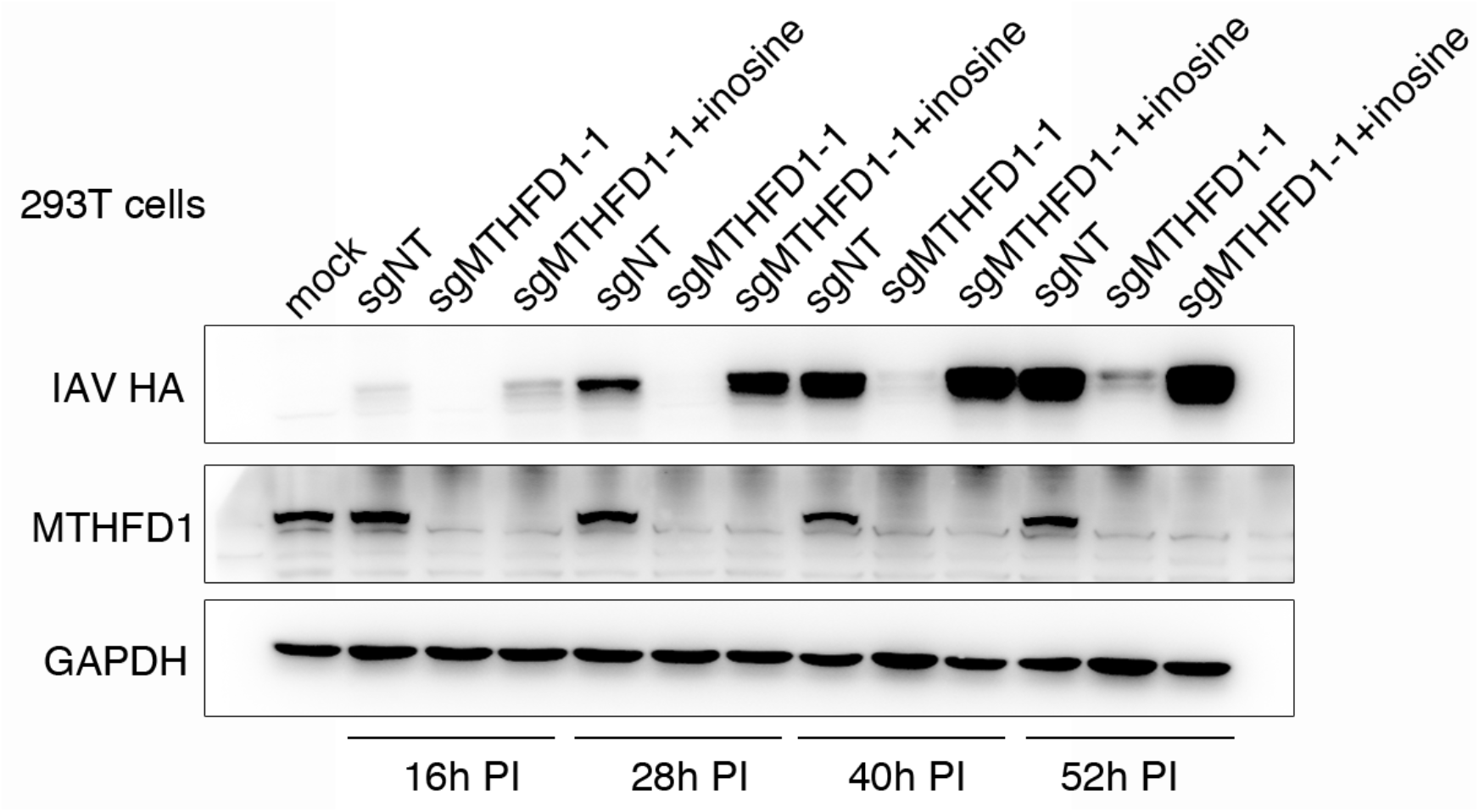
Supplementation of inosine rescues IAV replication in MTHFD1 knockout 293T cells. MTHFD1 knockout cells (sgMTHFD1-1) were starved in the medium without HT for 8h and then infected with PR8 with or without inosine supplementation. At 16h, 28h, 40h and 52h post infection, the cells were collected and analyzed by Western blotting.

**Supplementary Figure 5.**
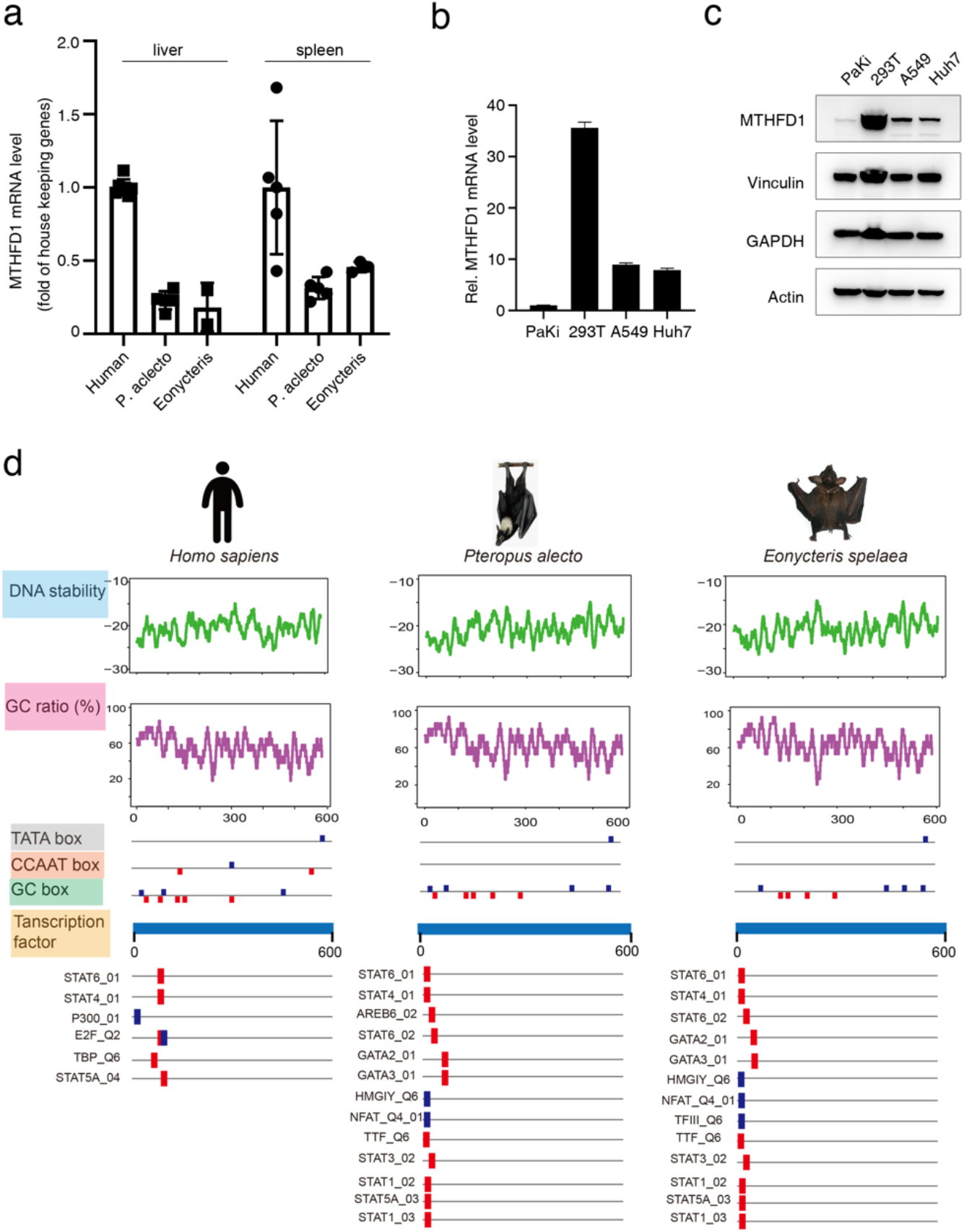
Bats have a lower MTHFD1 expression level compared humans. **a.** livers and spleens of human and two bat species were applied to RNAseq, and MTHFD1 mRNA level was analyzed. **b.** MTHFD1 mRNA levels in difference cell lines were compared using qRTPCR with GAPDH as a reference gene. **c.** MTHFD1 protein levels in different cell lines was compared by Western blotting. **d.** promoter analysis of MTHFD1 between human and two bat species *(P. alecto* and *E. spelaea).* The transcription factor binding sites and structure of promotors were characterized by the help of GPMiner (http://gpminer.mbc.nctu.edu.tw/) and the related information of the regulatory features such as transcription factor binding site (TFBS), G + C content, TATA box, CCAAT box, GC box and DNA stability are shown. Blue and red boxes indicate positions of those putative elements and related transcription factors predicted by GPMiner. A lack of CCAAT box was observed in both bats.

